# microRNA-138-5p suppresses excitatory synaptic strength at the cerebellar input layer

**DOI:** 10.1101/2024.09.24.613637

**Authors:** Igor Delvendahl, Reetu Daswani, Jochen Winterer, Pierre-Luc Germain, Nora Maria Uhr, Gerhard Schratt, Martin Müller

## Abstract

MicroRNAs are small, highly conserved non-coding RNAs that negatively regulate mRNA translation and stability. In the brain, microRNAs contribute to neuronal development, synaptogenesis, and synaptic plasticity. The microRNA 138-5p (miR-138-5p) controls inhibitory synaptic transmission in the hippocampus and is highly expressed in cerebellar excitatory neurons. However, its role in cerebellar synaptic transmission remains unkwown. Here, we investigated excitatory transmission within the cerebellum in mice expressing a sponge construct that sequesters endogenous miR-138-5p. Mossy fiber stimulation-evoked excitatory postsynaptic currents (EPSCs) in granule cells were significantly larger compared with controls. Furthermore, we observed larger miniature EPSC amplitudes, suggesting increased postsynaptic AMPA receptor numbers. High-frequency train stimulations revealed enhanced short-term depression following miR-138-5p downregulation. Together with computational modelling, this suggests a negative regulation of presynaptic release probability. Overall, our results demonstrate that miR-138-5p suppresses synaptic strength through pre- and postsynaptic mechanisms, providing a powerful mechanism for tuning excitatory synaptic input into the cerebellum.

## Introduction

Synaptic transmission underlies a broad variety of brain functions from sensory integration to learning and memory. The transmission efficacy of individual synapses is thus a key factor determining neural function. The efficacy or strength of synapses depends on their molecular composition and arrangement as well as their history of activity, and is tightly controlled to enable proper functioning of the nervous system (Clayton et al., 2024; Grant, 2012; Zoghbi and Bear, 2012). Synapses can adjust their transmission efficacy to allow desired, adaptive modifications (Bliss et al., 2013; Citri and Malenka, 2008), but also to counteract maladaptive deviations (Delvendahl and Müller, 2019). However, the mechanisms that allow such extensive and intricate control of synaptic strength are poorly understood.

The proteomic landscape of synapses is highly complex (Gonzalez-Lozano et al., 2020; Van Oostrum et al., 2023). Several key synaptic proteins have been attributed important functions for synaptic transmission or plasticity (Sheng and Kim, 2011; Südhof, 2012). Typically, the reduction or elimination of the expression of these proteins results in loss-of-function phenotypes, highlighting that many synaptic proteins facilitate synaptic transmission. Conversely, a subset of synaptic proteins acts to suppress synaptic strength by constraining neurotransmitter release and/or postsynaptic function, such as Tomosyn (McEwen et al., 2006), Mover (Körber et al., 2015), or SynGAP (Araki et al., 2015). Mechanisms that limit synaptic efficacy may be particularly important to maintain information processing and to prevent overexcitation of neural networks (Eiro et al., 2023; Leite et al., 2005). However, our understanding of the negative regulation of synaptic strength remains limited.

MicroRNAs (miRNAs) are small noncoding RNAs that can bind to the 3’ untranslated regions (UTRs) of multiple target mRNAs and suppress their translation (Filipowicz et al., 2008; Friedman et al., 2009). The biological functions of miRNAs are diverse. In the nervous system, miRNAs influence various processes ranging from neuronal development and synaptogenesis to memory and behavior (Brennan and Henshall, 2020; Konopka et al., 2010; Kosik, 2006; Soutschek and Schratt, 2023). At the synaptic level, specific miRNAs can control the abundance of postsynaptic AMPA receptors (AMPARs) (Hanley, 2021) and affect synaptic plasticity (Schratt, 2009). Certain miRNAs are enriched in synapto-dendritic compartments (Sambandan et al., 2017; Schratt et al., 2006; Siegel et al., 2009), suggesting an important role in the local regulation of synaptic function. Among these, miR-138-5p regulates the morphology of dendritic spines and has a suppressive influence on synaptic function in primary rat hippocampal neurons (Siegel et al., 2009). On a behavioral level, loss of miR-138-5p impairs short-term memory in mice, which is most likely due to an increase in inhibitory transmission in area CA1 of the hippocampus (Daswani et al., 2022). In hippocampal interneurons, miR-138-5p may modulate synaptic transmission by inhibiting expression of the schizophrenia susceptibility gene *Erbb4* (Erb-B2 receptor tyrosine kinase 4) (Daswani et al., 2022). miR-138-5p is strongly expressed in the cerebellar cortex (He et al., 2012; Obernosterer et al., 2006). In this brain region, miRNAs have been shown to be important during development (Constantin, 2017), but their functions in the adult cerebellum remain largely unexplored.

The cerebellum is involved in a wide range of tasks, from motor control to cognitive functions (Carey, 2024; De Zeeuw et al., 2021; Schmahmann, 2004). Its computational capabilities depend on a highly conserved, canonical circuit architecture (Albus, 1971; Apps et al., 2018; Marr, 1969). Excitatory inputs to the cerebellar cortex are provided by climbing fibers (CFs) and mossy fibers (MFs). CFs directly target Purkinje cells (PCs), whereas MFs primarily synapse with cerebellar granule cells (GCs). GCs are highly numerous and provide an increase in dimensionality and sparsening of sensory information, which is essential for pattern separation (Cayco-Gajic and Silver, 2019) and supports cerebellum-dependent learning (Kita et al., 2021). Information transfer at the MF–GC synapse is thus highly relevant for cerebellar function. However, the mechanisms controlling synaptic strength at this key cerebellar synapse are not well understood and very little is known about the negative regulation of MF–GC transmission.

Based on its prominent cerebellar expression, we hypothesized that miR-138-5p suppresses excitatory synaptic strength in the cerebellum. We quantified synaptic transmission at cerebellar MF–GC synapses in acute mouse brain slices upon expression of a sponge construct that causes a sequestering of miR-138-5p. Our findings reveal that synaptic strength is increased in miR-138 sponge-expressing GCs, driven by enhanced postsynaptic AMPAR function and elevated presynaptic release probability. Thus, miR-138-5p negatively regulates synaptic efficacy at MF–GC synapses via both presynaptic and postsynaptic mechanisms, providing powerful control over synaptic excitation at the cerebellar input layer.

## Results

### miR-138-5p targets synaptic genes and is expressed in cerebellar excitatory neurons

We generated mice with a conditional ROSA26 transgene (‘miR-138 floxed’), which allows the expression of a sponge transcript that inactivates endogenous miR-138-5p upon Cre-recombinase expression (Daswani et al., 2022). Sponge transcripts sequester endogenous miRNA, thereby leading to miRNA inactivation and the de-repression of cognate target genes (Ebert and Sharp, 2010). We activated miR-138-sponge expression at the embryonic stage by crossing 138-floxed mice to the ubiquitous Cre-driver line CMV-Cre (‘miR-138 sponge’). miR-138 floxed mice without CMV transgene served as controls. We first investigated if the miR-138 sponge construct is expressed in the cerebellum. lacZ staining revealed highly penetrant expression of the 6×-miR-138 sponge in the cerebellar cortex of miR-138 sponge mice (**Figure S1**).

To understand how miR-138-5p might regulate synaptic function in the cerebellum, we first examined the annotated functions of the strongest predicted miR-138-5p targets (i.e., 7/8mer) among genes expressed in cerebellar granule cells using over-representation of SynGO terms (Koopmans et al., 2019). This analysis revealed enrichments for genes that exert their function on presynaptic or postsynaptic compartments (**Figure 1A**). Given that synaptic genes tend to have a longer UTR, and as such are enriched for miRNA binding sites in general, we also established that these enrichments were much stronger for miR-138-5p targets than for most other miRNAs (**Figure 1A**). This indicates that miR-138-5p may regulate synaptic genes that are potentially related to pre- and postsynaptic function.

**Figure 1.**
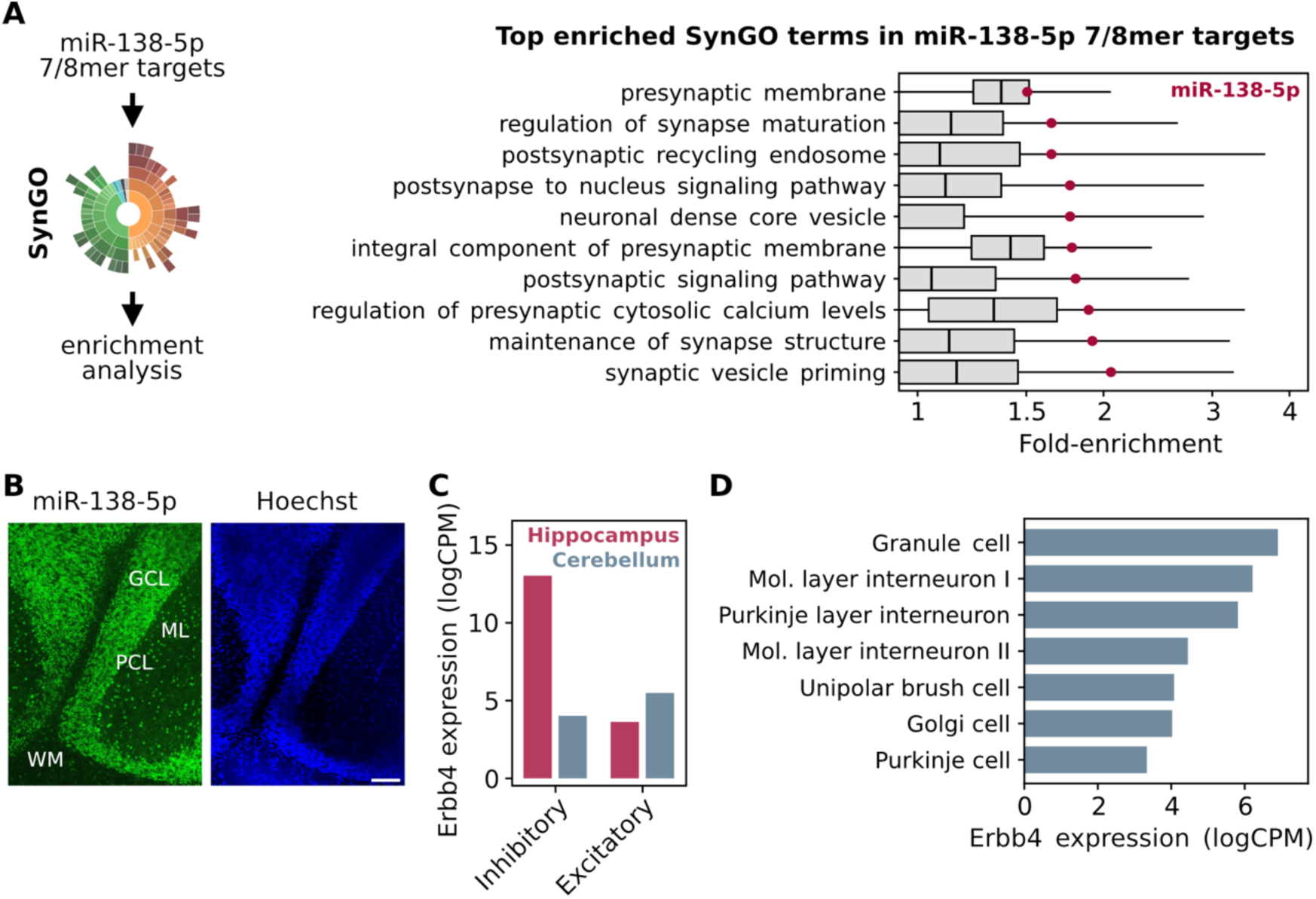
miR-138-5p targets synaptic genes and is expressed in cerebellar excitatory neurons. **(A)** Top 10 SynGO terms significantly enriched in the 7/8mer targets of miR-138-5p. Whereas the targets of many miRNAs are enriched for pre-/post-synaptic genes, miR-138-5p (red dots) is consistently amongst those showing strongest enrichment. **(B)** Single-molecule fluorescence in situ hybridization for miR-138-5p in the cerebellum (Left) and Hoechst nuclear staining (Right). Layers of the cerebellar cortex are indicated (GCL, granule cell layer; ML, molecular layer; PCL, Purkinje cell layer; WM, white matter). Scale bar, 100 µm. **(C)** The direct miR-138-5p target *Erbb4* (Daswani et al., 2022) is differentially expressed in inhibitory and excitatory neurons of the hippocampus and cerebellum. **(D)** Cerebellar expression of Erbb4, which has the highest expression levels in granule cells.

miR-138-5p is a brain-enriched miRNA with strong expression in the hippocampus and cerebellum (Obernosterer et al., 2006). We performed in-situ hybridization to further investigate the localization of miR-138-5p in the cerebellar cortex. Our results demonstrated strong expression of this miRNA in granule cells (**Figure 1B**), the most abundant excitatory neurons in the cerebellar cortex. Erbb4 is a direct miR-138-5p target (Daswani et al., 2022) that has been implicated in pre- and postsynaptic function (**Figure S1**). We analyzed publicly available single-cell RNAseq datasets to establish the expression levels of Erbb4 in the cerebellum and hippocampus. Our analysis revealed that the highest expression levels of Erbb4 can be detected in inhibitory hippocampal interneurons and excitatory cerebellar GCs, whereas it is barely expressed in excitatory neurons of the hippocampus (**Figure 1C–D**). At the synaptic level, Erbb4 is located at both postsynaptic and presynaptic compartments (Koopmans et al., 2019). Together, these results show that miR-138-5p is highly expressed in the adult cerebellar cortex. In addition, they suggest that miR-138-5p is a prominent candidate capable of modulating synaptic properties on both pre- and postsynaptic sides, and that cerebellar GCs highly express Erbb4, a direct target of miR-138-5p.

### miR-138-5p negatively regulates synaptic strength at cerebellar MF–GC synapses

To determine the role of miR-138-5p in excitatory neurotransmission within the cerebellum, we performed whole-cell patch-clamp recordings from GCs in acute slices from adult mice. We isolated AMPAR-mediated synaptic transmission and recorded excitatory postsynaptic currents (EPSCs) from GCs of control (miR-138 floxed) and miR-138 sponge mice. Stimulation of MFs resulted in considerably larger evoked EPSCs in sponge GCs, with ~37% bigger amplitudes than in control synapses (**Figure 2A–D**). This increase in EPSC amplitude due to miR-138-5p inactivation was consistent across individual animals (**Figure S2**). Furthermore, EPSC charge transfer was increased in miR-138 sponge GCs (**Figure 2E**), and we noted a slight slowing of the EPSC decay without changes in the risetime (**Figure 2F–G**). These findings indicate that miR-138-5p negatively regulates synaptic efficacy at cerebellar MF–GC synapses by modulating AMPAR-mediated transmission. Interestingly, synaptic strength at parallel fiber to Purkinje cell synapses was not enhanced in miR-138 sponge mice (**Figure S3**), pointing towards a synapse-specific regulatory role of this miRNA within the cerebellum.

**Figure 2.**
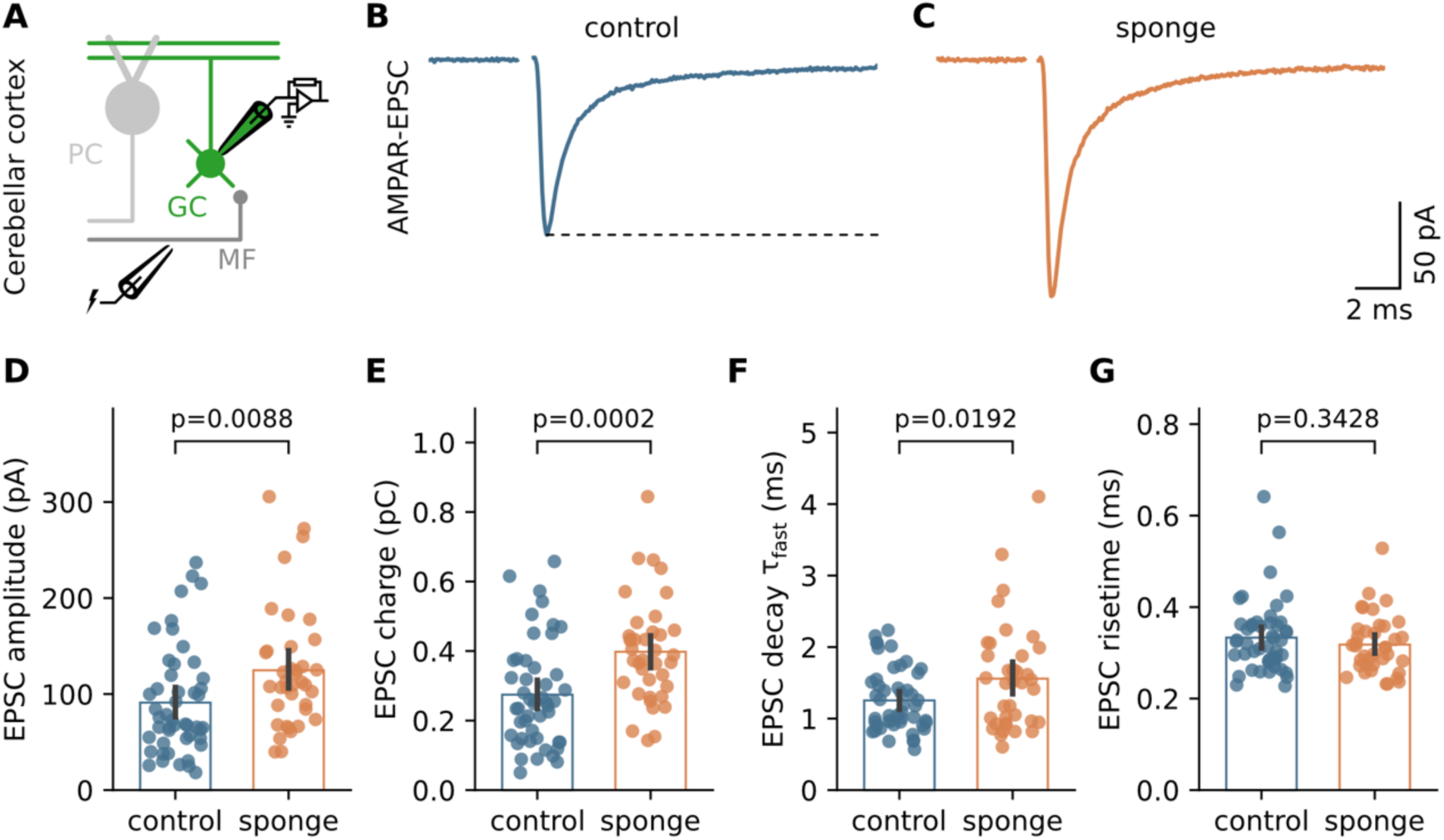
miR-138-5p negatively regulates synaptic strength at cerebellar MF–GC synapses. **(A)** Cartoon illustrating MF–GC recordings in acute cerebellar slices from adult control and miR-138 sponge mice. **(B)** Example average EPSC recorded from a control MF–GC synapse. Average of 24 sweeps; stimulation artifact is blanked. **(C)** Example EPSC from a miR-138 sponge synapse (average of 18 sweeps). **(D)** Quantification of EPSC amplitude for control and sponge conditions. **(E)** miR-138 sponge MF–GC EPSCs have increased charge. **(F)** The fast EPSC decay time constant is slightly slower in miR-138 sponge GCs. **(G)** EPSC 10–90% risetime is similar for both genotypes.

### miR-138-5p negatively regulates AMPAR numbers in cerebellar GCs

The enhanced synaptic efficacy observed following downregulation of miR-138-5p could be attributed to an increase in AMPAR function or number. To probe if AMPARs are involved in the EPSC amplitude increase, we recorded spontaneous miniature EPSCs (mEPSCs) from cerebellar GCs. GCs from miR-138 sponge mice exhibited increased mEPSC amplitudes (**Figure 3A–C**, **Figure S4**), indicating enhanced AMPAR-mediated transmission. This finding aligns with previous reports of reduced mEPSC amplitudes upon miR-138-5p overexpression in hippocampal cultures (Siegel et al., 2009). Additionally, mEPSC charge was slightly increased in sponge GCs (**Figure 3D**), whereas the decay kinetics and frequency of mEPSCs were similar across both genotypes (**Figure 3E–F**). mEPSC amplitude mainly depends on synaptic AMPAR numbers and conductance. To determine whether miR-138-5p influences AMPAR single-channel conductance or receptor numbers, we performed fluctuation analysis of the mEPSC decay (Traynelis et al., 1993) (**Figure 3G–H**). Single-channel conductance was comparable between conditions (control: 14.1 pS, sponge: 14.0 pS, **Figure 3I**), but the number of channels was greater in sponge GCs (**Figure 3J**). This suggests that miR-138-5p negatively regulates the number of synaptic AMPARs in cerebellar GCs. Together, mEPSC analysis revealed larger mEPSC amplitudes due to an increased number of postsynaptic AMPARs in miR-138 sponge GCs, consistent with the idea that miR-138 negatively regulates synaptic AMPAR levels.

**Figure 3.**
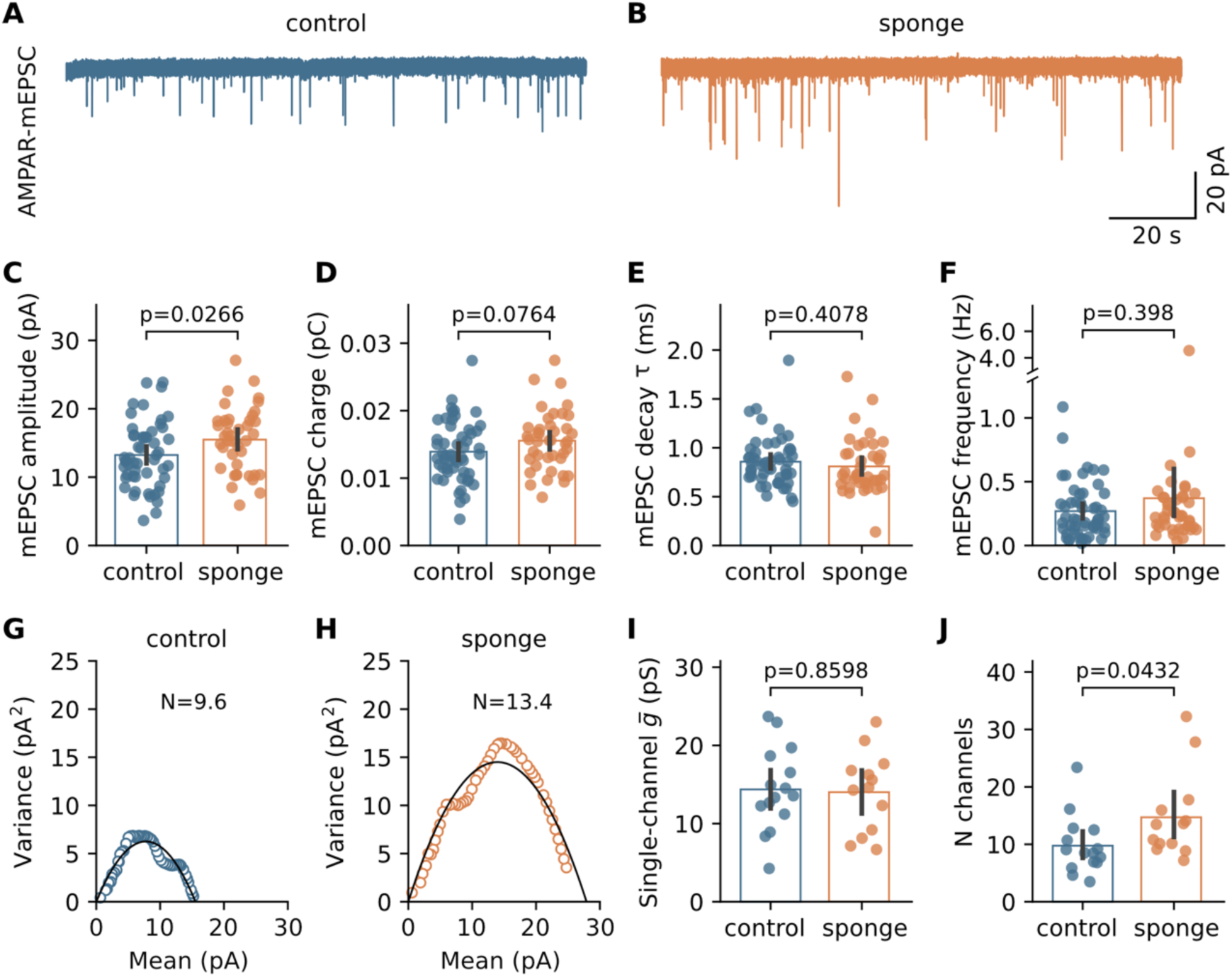
miR-138-5p negatively regulates AMPAR numbers in cerebellar GCs. **(A)** Example mEPSC recording from a control GC. **(B)** Example recording from a miR-138 sponge GC. **(C)** Mean mEPSC amplitude is bigger in sponge than in control GCs. **(D)** Quantification of mEPSC charge for both conditions. **(E)** Quantification of mEPSC decay time constant, calculated from exponential fits to the average mEPSC waveform of each cell. **(F)** The frequency of mEPSCs is similar in both genotypes. **(G–H)** Peak-scaled mEPSC decay variance versus mean amplitude for both genotypes. **(I)** AMPAR single-channel conductance was unaltered by miR-138 sponge expression. **(J)** AMPAR number is higher in sponge GCs.

### miR-138-5p suppresses presynaptic function

Interestingly, the mEPSC amplitude increase only accounts for about half of the overall increase in synaptic efficacy following miR-138 sponge expression (17.0% vs. 36.9%, Cohen’s d 0.48 [95%CI 0.06, 0.90] vs. 0.69 [95%CI 0.26, 1.11]), suggesting that the role of miR-138-5p at MF–GC synapses extends beyond simply regulating postsynaptic AMPAR levels. To investigate if presynaptic mechanisms contribute to the enhanced synaptic efficacy at miR-138 sponge MF–GC synapses, we analyzed quantal content – a proxy for neurotransmitter release – derived from evoked EPSCs and mEPSC recordings in the same cells. We found that quantal content was increased in miR-138 sponge GCs (**Figure 4A**), indicating enhanced presynaptic glutamate release from MFs. Further analysis of evoked EPSC variance revealed a decreased coefficient of variance (CV), which is inversely correlated with presynaptic release probability, following downregulation of miR-138-5p (**Figure 4B–D**), suggesting an increased release probability. To further examine changes in release probability, we recorded EPSC paired-pulse ratios (PPRs, **Figure 4E–F**). MF–GC synapses show a combination of synaptic facilitation and depression upon paired-pulse stimulation, with depression prevailing at shorter intervals (Saviane and Silver, 2006). PPRs were reduced in miR-138 sponge GCs (**Figure 4G–H**), consistent with an increased release probability. A reduction in PPR could also arise from altered receptor desensitization. To test this possibility, we fit the relationship between PPR and inter-stimulus interval with bi-exponential fits (**Figure 4I**). The fast time constants were comparable for both conditions (control: 11.1 ms, sponge: 11.7 ms), suggesting a similar time course of AMPAR desensitization (DiGregorio et al., 2007; Saviane and Silver, 2006), therefore further supporting an increase in release probability in miR-138 sponge MF–GC synapses. PPRs of individual connections were negatively correlated with synaptic strength at control synapses (R = −0.58, **Figure 4J**). Interestingly, we only rarely observed facilitating MF–GC synapses in miR-138 sponge mice (**Figure 4K**, **Figure S5**), resulting in a weaker correlation between PPR and EPSC amplitude (R = −0.22, **Figure 4K**). In summary, miR-138 sponge synapses exhibit increased glutamate release and reduced PPR, indicating that miR-138-5p negatively regulates presynaptic release probability.

**Figure 4.**
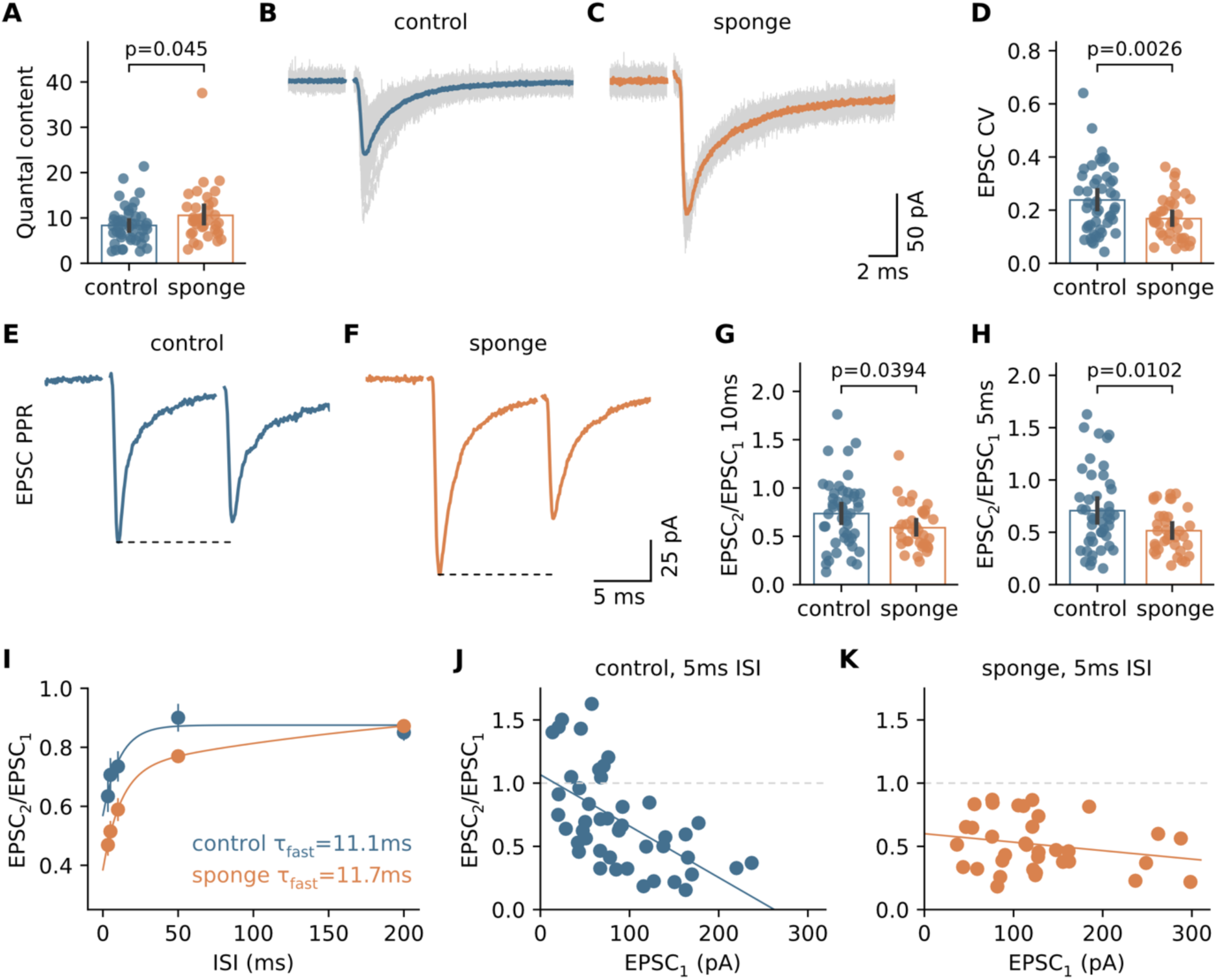
miR-138-5p suppresses presynaptic function. **(A)** Quantification of quantal content (=EPSC/mEPSC) for both genotypes. **(B)** Example individual EPSCs (gray) overlaid with the average (blue) for a control MF–GC synapse. Stimulation artifacts are blanked. **(C)** Example EPSCs from a sponge synapse overlaid with average (orange). **(D)** Coefficient of variation (CV) of evoked EPSC amplitudes is decreased in miR-138 sponge GCs. **(E)** Example paired-pulse responses from a control cerebellar GC (average of 5 consecutive sweeps). Stimulation artifacts are blanked. **(F)** Example recording from a sponge GC. **(G)** Paired-pulse ratios (PPRs, i.e., EPSC_2_/EPSC_1_) at 100 Hz for control and sponge GCs. **(H)** Same as in **G**, but for 200 Hz. **(I)** PPR versus inter-stimulus interval (ISI). Lines are bi-exponential fits, time constant of the fast component is indicated. **(J)** EPSC PPR versus amplitude of the first EPSC in GCs of control mice for 200 Hz stimulation with linear regression. **(K)** Same as (**J**), but for miR-138 sponge GCs.

### miR-138-5p reduces short-term depression at MF–GC synapses

To further analyze the effect of miR-138-5p downregulation on release probability and synaptic short-term plasticity, we performed high-frequency train stimulation experiments. MF–GC synapses are capable of operating with a high bandwidth of presynaptic action potential frequencies (Delvendahl and Hallermann, 2016), and typically exhibit prominent short-term depression upon sustained stimulation (Saviane and Silver, 2006). We recorded short trains of EPSCs at 300 Hz (**Figure 5A–B**), followed by single pulses to monitor recovery from depression (Hallermann et al., 2010). In miR-138 sponge GCs, we observed increased synaptic depression (**Figure 5C**), evident from a faster time constant of depression (**Figure 5D**) and reduced steady-state amplitudes (**Figure 5E**). This more pronounced depression indicates an increase in release probability. To estimate the size of the readily releasable pool (RRP) of synaptic vesicles and apparent release probability, we analyzed cumulative EPSC amplitudes (Schneggenburger et al., 1999) (**Figure 5F**). RRP size was similar between control and miR-138 sponge synapses (**Figure 5G**), whereas release probability was increased in miR-138 sponge animals (Cohen’s d: 0.44 [95%CI −0.07, 0.94], **Figure 5H**). The steady-state recruitment slope was comparable between both genotypes (**Figure S6**). These EPSC high-frequency train recordings demonstrate altered synaptic short-term dynamics, with a stronger degree of depression in miR-138 sponge mice. Collectively, the results from paired-pulse EPSCs and train recordings support the notion that miR-138-5p negatively regulates release probability and short-term depression at cerebellar MF–GC synapses.

**Figure 5.**
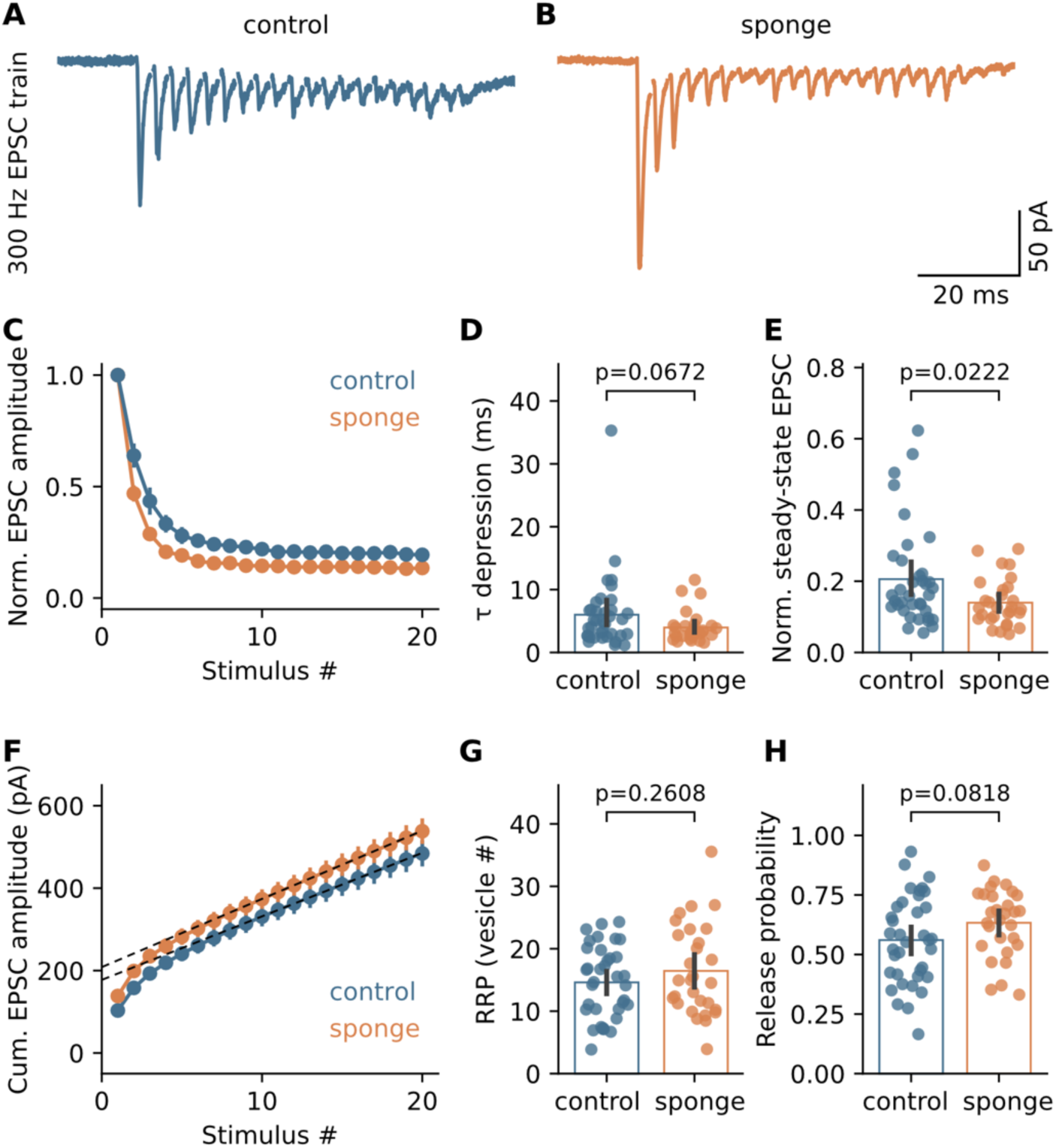
miR-138-5p reduces short-term depression at MF–GC synapses. **(A)** Example EPSC responses upon 300-Hz stimulation for a control GC. Average of five sweeps; stimulation artifacts are blanked for clarity. **(B)** Example 300-Hz train for a miR-138 sponge synapse. **(C)** Normalized EPSC amplitude for both genotypes. Error bars are SEM. **(D)** Time constant of synaptic depression for control and sponge synapses. **(E)** miR-138 sponge expression decreases the normalized steady-state EPSC amplitude. **(F)** Cumulative EPSC amplitude for 300-Hz train stimulation. Error bars represent SEM, dashed lines are linear fits to the last 10 datapoints. **(G)** Readily releasable pool (RRP) of synaptic vesicles, calculated from cumulative EPSC analysis. **(H)** Apparent release probability (first EPSC/cumulative EPSC) is higher in miR-138 sponge GCs.

### miR-138-5p negatively regulates presynaptic release probability

To investigate if increased release probability accounts for the observed short-term depression effects in miR-138 sponge GCs, we we applied a synaptic short-term plasticity model (Tsodyks et al., 1998) to our experimental data (**Figure 6A–B**). Across recordings, the fitted model parameter ‘utilization of synaptic efficacy’ (U_SE_), which corresponds to release probability, was elevated in the sponge condition (**Figure 6C**). These computational modelling results are consistent with a higher release probability driving the more pronounced short-term depression upon high-frequency stimulation in miR-138 sponge expressing synapses. In addition, the fitted model indicated a faster recovery time constant for the miR-138 sponge condition (**Figure S7**), pointing towards faster recovery from depression. We further analyzed the recovery from synaptic depression following 300-Hz stimulation in control and sponge GCs (**Figure S7**). Recovery at MF–GC synapses follows a bi-exponential time course (Hallermann et al., 2010). Consistent with the modelling results, the fast component of EPSC recovery was enhanced in sponge GCs, with a faster time constant and larger fractional amplitude (**Figure S7**). Thus, the increased synaptic depression in miR-138 sponge synapses is accompanied by faster recovery. Together, these findings strongly suggest that miR-138-5p suppresses release probability at MF–GC synapses.

**Figure 6.**
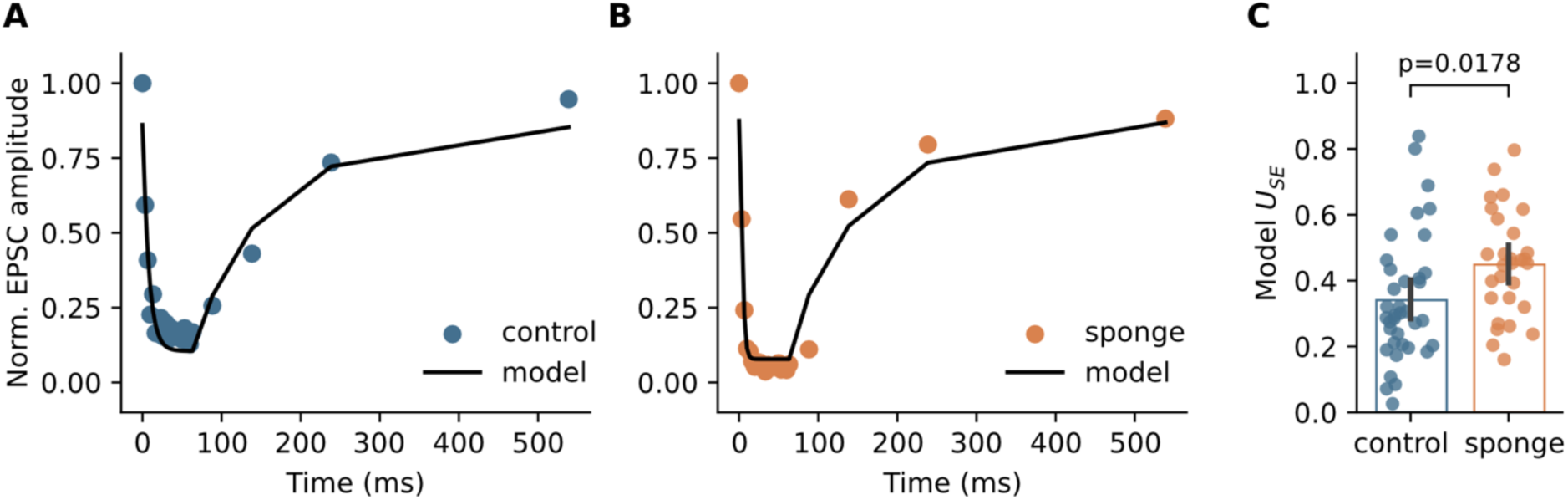
miR-138-5p negatively regulates presynaptic release probability. **(A)** Example normalized EPSC amplitudes for 300-Hz train stimulation and recovery pulses. Black line represents a Tsodyks-Markram short-term plasticity model fit to the experimental data. **(B)** Same as in **A**, but for an example miR-138 sponge synapse. **(C)** Quantification of model parameter U_SE_.

## Discussion

In this study, we revealed a general repressive mechanism for excitatory synaptic strength mediated by miR-138-5p at excitatory mossy fiber to granule cell synapses in the mammalian cerebellum. The expression of a miR-138-5p sponge construct caused an increase in postsynaptic mEPSC size via an increase in AMPAR numbers at synapses. In addition, sequestering of miR-138-5p led to enhanced glutamate release, driven by an increase in presynaptic release probability. miR-138-5p thus suppresses synaptic strength at excitatory cerebellar synapses through distinct presynaptic and postsynaptic mechanisms. This dual regulatory role underscores the importance of miR-138-5p in modulating synaptic excitation at the cerebellar input layer.

The distinct modulations at both sides of the synapse indicate that the effect of miR-138-5p on excitatory synaptic transmission extends beyond a simple regulation of an individual target mRNA. miR-138-5p rather acts as a general repressor of synaptic strength. Many different proteins influence the functional properties of individual synapses (Cizeron et al., 2020; Van Oostrum et al., 2023; Zhu et al., 2018). An extended line of research has investigated the role of individual proteins for synaptic function. Loss or downregulation of a given synaptic protein often caused a disruption of synaptic function and hence reduced synaptic strength, suggesting important roles of individual proteins for promoting synaptic transmission. By contrast, much less is known about the negative regulation of synaptic transmission. Our data provide evidence for a general, negative regulation of excitatory synaptic strength at cerebellar input synapses. A mechanism that suppresses synaptic strength under baseline conditions could enable dynamic modulation of synaptic function, such as activity-dependent (Mapelli et al., 2015) or homeostatic plasticity (Delvendahl and Müller, 2019). A miRNA-dependent mechanism controlling synaptic strength is thus an intriguing candidate for adaptive plasticity of synaptic function. Interestingly, miR-138 has been linked to memory formation in rodents (Li et al., 2018) and memory performance in humans (Schröder et al., 2014).

Our data point towards a brain region-specific, and potentially synapse type-specific, role of miR-138-5p for synaptic transmission. In the hippocampus, inactivation of miR-138-5p selectively affects inhibitory transmission onto pyramidal neurons (Daswani et al., 2022). In the cerebellum, we here identified a strong impact of miR-138 sponge expression on excitatory MF–GC synapses. The negative regulation of excitatory synaptic strength may be specific to MF–GC synapses, because transmission at parallel fiber synapses onto Purkinje cells – which also express miR-138-5p (Zolboot et al., 2023) – was not increased. It will be interesting to further elucidate the function of miR-138-5p in Purkinje cells, and to study if this miRNA has effects on inhibitory synapses in the cerebellum. A presumably synapse type-specific regulatory effect suggests that miRNAs like miR-138-5p could fine-tune synaptic properties in a circuit-specific manner.

Overexpression of miR-138-5p decreased mEPSC amplitudes in a hippocampal culture preparation (Siegel et al., 2009). In combination with our observation of enhanced mEPSC amplitudes upon miR-138 sponge expression in a cerebellar slice preparation, these findings strongly suggest that miR-138-5p negatively regulates synaptic AMPAR levels. A previous study identified Erbb4 as a direct target of miR-138-5p (Daswani et al., 2022). Erbb4 is a receptor tyrosine kinase that can interact with postsynaptic density proteins (Garcia et al., 2000). Erbb4 has a prominent role in inhibitory hippocampal interneurons, where miR-138 sponge expression caused an increase in inhibitory miniature event frequency (Daswani et al., 2022). The excitatory GCs in the cerebellar cortex also express Erbb4, suggesting a region-specific role of this receptor tyrosine kinase. Interestingly, Erbb4 was shown to regulate translation of several synaptic proteins, including the pore-forming AMPAR subunit GluA4 (Bernard et al., 2022). In the cerebellum, Erbb4 co-precipitates with GluA4 (Pelkey et al., 2015), which is the predominant AMPAR subunit in GCs (Kita et al., 2021). In our study, inactivation of miR-138-5p increased mEPSC amplitudes in cerebellar GCs. This increase is likely mediated by increasing synaptic AMPAR numbers, which would be consistent with miR-138-5p targeting (and thus repressing) Erbb4, which in turn promotes GluA4 translation. A regulation of AMPAR subunits by miRNAs has been demonstrated for GluA1 and GluA2. miR-183/96 double KO increased GluA1 levels at the calyx of Held synapse (Krohs et al., 2021) and GluA1 is also a direct target of miR-137 (Olde Loohuis et al., 2015) and miR-92a (Letellier et al., 2014). Likewise, miR-186 (Silva et al., 2019), miR-124 (Ho et al., 2014), as well as miR-181 (Saba et al., 2012) regulate GluA2 levels in hippocampal cultures. miRNA-dependent regulation of AMPAR levels may thus be a pervasive mechanism of controlling excitatory synaptic strength (Hanley, 2021).

In addition to postsynaptic effects on the level of AMPAR function, miR-138-5p also prominently affected presynaptic function at cerebellar MF–GC synapses. The combined presynaptic and postsynaptic effect on synaptic efficacy is reminiscent of results on another miRNA complex – miR-183/96 – at the calyx of Held synapse in the auditory brainstem (Krohs et al., 2021). In contrast to our observation of a miR-138-5p-dependent regulation of release probability at cerebellar MF–GC synapses, global knock-out of miR-183 and miR-96 increased RRP size without affecting release probability at the calyx of Held. Nonetheless, the convergent negative regulation of both synaptic compartments by distinct miRNAs in different brain regions points towards a general mechanism for controlling excitatory synaptic strength. This notion is further supported by data from the *Drosophila* neuromuscular junction, where miR-34 controls synaptogenesis by regulating distinct pre- and postsynaptic genes (McNeill et al., 2020).

miRNAs can potentially influence release probability by repressing the translation of presynaptic proteins that are involved in setting release probability. While miR-138-5p has some predicted targets with presynaptic localization, experimental evidence for its regulation of presynaptic targets is currently lacking. The direct miR-138-5p target Erbb4 (Daswani et al., 2022) is linked to both presynaptic and postsynaptic compartments (Koopmans et al., 2019). Intriguingly, Erbb4 may also promote presynaptic release probability (Wang et al., 2018) and could thus be involved in the altered release properties that we observed in miR-138 sponge mice. Nevertheless, it remains unclear if Erbb4 is also a direct target of miR-138-5p in the cerebellum, and if the effects of miR-138-5p inactivation on presynaptic function are due to expression of this miRNA in presynaptic cells or due to trans-synaptic signalling. The negative regulation of presynaptic function by miR-138-5p is also consistent with findings in *Drosophila*, where another miRNA, miR-130, was found to suppress quantal content via negative regulation of the active zone protein Bruchpilot and Ca^2+^ influx (Tsurudome et al., 2010). A negative regulation of active zone proteins was also shown for miR-153 and miR-137 in the mouse hippocampus (Mathew et al., 2016; Siegert et al., 2015). miRNAs may thus control the levels of key active zone proteins, thereby regulating presynaptic functions such as release probability.

The influence of miR-138-5p on presynaptic release appears to enable low release probability synapses with facilitating short-term dynamics. This effectively broadens the distribution of synaptic strengths and short-term plasticity behavior across the population of MF–GC synapses, which may have important implications for cerebellar network function. For example, diversity of synaptic short-term dynamics at the cerebellar input layer could serve as a mechanism for temporal learning (Barri et al., 2022). A broad distribution of MF– GC short-term plasticity may also enhance pattern separation (Chabrol et al., 2015), which is thought to be a key computation of the cerebellar cortex. Our data indicate that miR-138-5p contributes to synaptic diversity at the cerebellar input layer and might support cerebellar computations by endowing a fraction of MF–GC synapses with facilitating properties. It will be highly interesting to study if inactivation of miR-138-5p affects cerebellum-dependent behavior or learning.

In summary, we investigated the role of miR-138-5p for excitatory synaptic transmission in the mouse cerebellum. Expressing an inactivating sponge construct of this miRNA prominently increased synaptic efficacy at MF–GC synapses through combined presynaptic and postsynaptic mechanisms. We conclude that miR-138-5p acts as a negative regulator of synaptic strength at the cerebellar input layer. The general repression of synaptic efficacy across synaptic compartments provides a powerful regulatory mechanism for synaptic function, which may be crucial for maintaining proper cerebellar network function.

## Methods

### Animals

Experiments were performed in C57BL/6NTac-Gt(ROSA)26So^tm2459(LacZ, antimir_138)Arte^ (‘miR-138 floxed’) and 138-sponge^ub^ (‘mir-138 sponge’) mice. Details of the generation of these animals have been reported previously (Daswani et al., 2022). Animals were treated in accordance with national and institutional guidelines. All experiments were approved by the Cantonal Veterinary Office of Zurich (authorization no. ZH206/16 and ZH194/21). Mice were housed in groups of 3–5 per cage, with food and water ad libitum. Recordings were performed in adult (4–8 months old) male mice using four miR-138 sponge animals and five littermate miR-138-floxed controls.

### LacZ staining

β-galactosidase activity was detected using the chromogenic substrate X-gal. Fresh frozen brain sections were fixed with formalin and after washing incubated in X-gal working solution at 37 °C for 24 h. After incubation, slides were washed and mounted for imaging (as described in https://ihcworld.com/2024/01/26/x-gal-staining-protocol-for-beta-galactosidase/).

### Single-molecule fluorescence in situ hybridization (smFISH)

smFISH for miRNA detection on cerebellar slices was performed using the ViewRNA Tissue Assay Fluorescence Kit (Thermo Fisher) with miRNA pretreatment. The protocol followed the manufacturer’s instructions, with two modifications: an additional baking step of 13 minutes at 60 °C, and a reduced protease treatment duration of seven minutes. 16-µm thick cerebellar slices were prepared using a cryostat (Leica). Brain tissues were collected after perfusion of the animal with 4% paraformaldehyde (PFA), followed by a gradual increase in sucrose concentration to 30% over three days. The slices were stored in OCT at –80 °C until sectioning. Images were acquired with a confocal laser scanning microscope (CSLM 880, Zeiss) at 10× magnification.

### Gene expression data

For gene expression in the cerebellum, we used the single-nuclei RNA-seq data from Kozareva et al. (Kozareva et al., 2021) (accession id: GSE165371), specifically using the P60 mice samples. For gene expression in the hippocampus, we used the single-nuclei RNA-seq data from von Ziegler et al. (Von Ziegler et al., 2022) (accession id: GSE169510). In both cases, we produced pseudobulk samples from the quantification and cell annotations provided by the authors.

### miRNA target prediction and enrichment analysis

To predict miRNA targets, we took the longest annotated UTR of each protein-coding gene in the Ensembl GRCm38 release 99 annotation, and scanned it for 7/8mer canonical sites using scanMiR (Soutschek et al., 2022). For enrichment analysis, we used the SynGO release 20231201 (Koopmans et al., 2019), and used genes expressed at >2 log counts-per-million (CPM) in granule cells as a background. We excluded genes without any SynGO annotation from the background. For each miRNA, we tested the 7/8mer predicted targets for over-representation (hypergeometric test) of all SynGO terms that had at least 10 annotated genes in the filtered universe. Reported are the top 10 most significant terms for miR-138-5p targets, all with FDR below computing accuracy.

### Electrophysiology

Mice were sacrificed by rapid decapitation according to national guidelines. The cerebellar vermis was removed quickly and mounted using superglue in a chamber filled with cooled extracellular solution. Parasagittal 300-µm-thick slices were cut using a Leica VT1200S vibratome (Leica Microsystems, Germany), transferred to an incubation chamber at 35 °C for 30 min and then stored at room temperature until experiments. The extracellular solution (artificial cerebrospinal fluid, ACSF) for slice cutting and storage contained (in mM): 125 NaCl, 25 NaHCO_3_, 20 D-glucose, 2.5 KCl, 2 CaCl_2_, 1.25 NaH_2_PO_4_, 1 MgCl_2_, aerated with 95% O_2_, and 5% CO_2_.

Cerebellar slices were visualized using an upright microscope equipped with a 60×, 1 NA water immersion objective, infrared optics, and differential interference contrast (Scientifica, UK). The recording chamber was continuously perfused with ACSF supplemented with 10 µM D-APV, 10 µM bicuculline, and 1 µM strychnine to isolate AMPAR-mediated transmission. Voltage-clamp recordings were done using a HEKA EPC10 amplifier (HEKA Elektronik GmbH, Germany). Data were filtered at 10 kHz and digitized with 200 kHz; recordings of spontaneous postsynaptic currents were filtered at 2.7 kHz and digitized with 50 kHz. All experiments were performed at room temperature (22–25 °C). Patch pipettes were pulled to open-tip resistances of 5–8 MΩ (when filled with intracellular solution) from borosilicate glass (Science Products, Germany) using a DMZ puller (Zeitz Instruments, Germany). The intracellular solution contained (in mM): 150 K-D-gluconate, 10 NaCl, 10 HEPES, 3 MgATP, 0.3 NaGTP, 0.05 ethyleneglycol-bis(2-aminoethylether)-N,N,N’,N’-tetraacetic acid (EGTA), pH adjusted to 7.3 using KOH. Voltages were corrected for a liquid junction potential of +13 mV.

Electrophysiological recordings were performed blinded to genotype. We recorded from visually identified GCs in lobules III–VI of the cerebellar vermis. Spontaneous miniature excitatory postsynaptic currents were recorded at a holding potential of –100 mV with typical recording duration of 120–360 s. Extracellular MF stimulation was performed using bipolar square voltage pulses (duration, 150 µs) generated by an ISO-STIM 01B stimulus isolation unit (NPI, Germany) and applied through an ACSF-filled pipette. The pipette was moved over the slice surface in the vicinity of the patched GC while applying voltage pulses until excitatory postsynaptic currents (EPSCs) could be evoked reliably. Care was taken to stimulate single mossy fiber inputs, as demonstrated by robust average EPSC amplitudes when increasing stimulation intensity (Silver et al., 1996). EPSC recordings were performed at a frequency of 0.1 Hz with stimulation intensity 1–2 V above the threshold, typically <20 V. For high-frequency train stimulation, we applied 20 pulses at 100 or 300 Hz, followed by six single pulses to monitor recovery from depression. Recovery pulses were spaced at inter-stimulus intervals of 25, 50, 100, 300, 1000 and 3000 ms. Train stimulations were repeated five times with 30 s interval.

### Data analysis

Data analysis was performed blinded to genotype. Spontaneous mEPSCs were detected and analyzed using *miniML* (O’Neill et al., 2024) with a model trained on cerebellar GC mEPSCs. Evoked EPSCs were analyzed using custom-written routines in IgorPro (WaveMetrics). EPSC amplitudes were measured as peak-to-baseline following 50-point binomial smoothing of the recordings. Baseline was averaged from a 2-ms window before stimulation, and peaks were calculated from a 100-µs window centered around the EPSC minimum. For train-stimulation EPSCs, a 400-µs baseline window after the stimulation artifact was used. EPSC charge was integrated over the entire duration of the EPSC, and decay times were extracted from biexponential fits to the decay of the averaged EPSC of each cell. AMPA receptor single-channel conductance was analyzed from the variance of peak-scaled mEPSC decays (Traynelis et al., 1993) using recordings with >50 detected events as previously described (Delvendahl et al., 2019). For calculation of quantal content and RRP, mEPSC amplitudes were scaled according to the difference in holding voltage (–100 mV vs. –80 mV), assuming an AMPAR current reversal potential of 0 mV. Recovery from short-term depression was analyzed normalized to the first EPSC amplitude of the train stimulation. To quantify the time course of recovery, we fit normalized EPSC amplitudes with a bi-exponential function. We only included synapses that reached at least 85% of the initial EPSC amplitude in the ~5 s interval covered by the single recovery EPSC stimulations. Traces shown in figures were filtered using a 15–25-samples Hann window for display purposes.

### Computational modelling

We fit a Tsodyks-Markram model (Tsodyks et al., 1998; Tsodyks and Markram, 1997) to the MF–GC EPSC train data to extract the utilization of synaptic efficacy (U_SE_). This model parameter represents the fraction of (presynaptic) resources used upon an action potential. The model parameters (U_SE_, A_SE_, ρ_facilitation_, ρ_recovery_) were optimized for each recording using normalized EPSC amplitudes recorded at 100 and 300 Hz. We ran model optimization via an evolutionary algorithm (Blue Brain Python Optimisation Library) with offspring size 500, η of 20, mutation probability 0.4 and crossover probability 0.7. Simulations were run for a maximum of 50 generations. Simulations were implemented in Python 3.9 using the *NEURON* (Hines, 2009) and *BluePyOpt* (Van Geit et al., 2016) libraries.

### Statistics

We used the Python library *dabest* for statistical analyses (Ho et al., 2019). Effect sizes are reported as Cohen’s d with 95% confidence intervals, obtained by bootstrapping (5000 samples; the confidence interval is bias-corrected and accelerated) (Ho et al., 2019). Statistical comparisons were performed using permutation t-tests with 5000 reshuffles.

## Data Availability Statement

All data supporting the results in the manuscript are included within the figures and/or text. Raw data is available from the corresponding author upon reasonable request.

## Acknowledgments

We thank Yannick Ruhl and Jochen Schwenk for critical reading of the manuscript.

## Funding statement

This work was supported by the Swiss National Science Foundation (grants PZ00P3_174018 to I.D. and PP00P3_144816 to M.M.), the German Research Foundation (grant no. 535029399 to I.D.), a European Research Council Starting Grant (SynDegrade; grant 679881 to M.M.), and a University Research Priority Program “Adaptive Brain Circuits in Development and Learning” grant by the University of Zurich (to M.M.).

## Author contributions

**I.D.:** Conceptualization, Investigation, Formal analysis, Visualization, Writing - Original Draft, Funding acquisition. **R.D.:** Investigation, Formal analysis. **J.W.:** Conceptualization, Formal analysis, Writing - Review & Editing. **P.-L.G.:** Investigation, Formal analysis, Writing - Review & Editing. **N.U.:** Investigation, Formal analysis. **G.S.:** Conceptualization, Supervision, Funding acquisition, Writing - Review & Editing. **M.M.:** Conceptualization, Supervision, Funding acquisition, Writing - Review & Editing.

## Supplementary Figures

**Figure S1.**
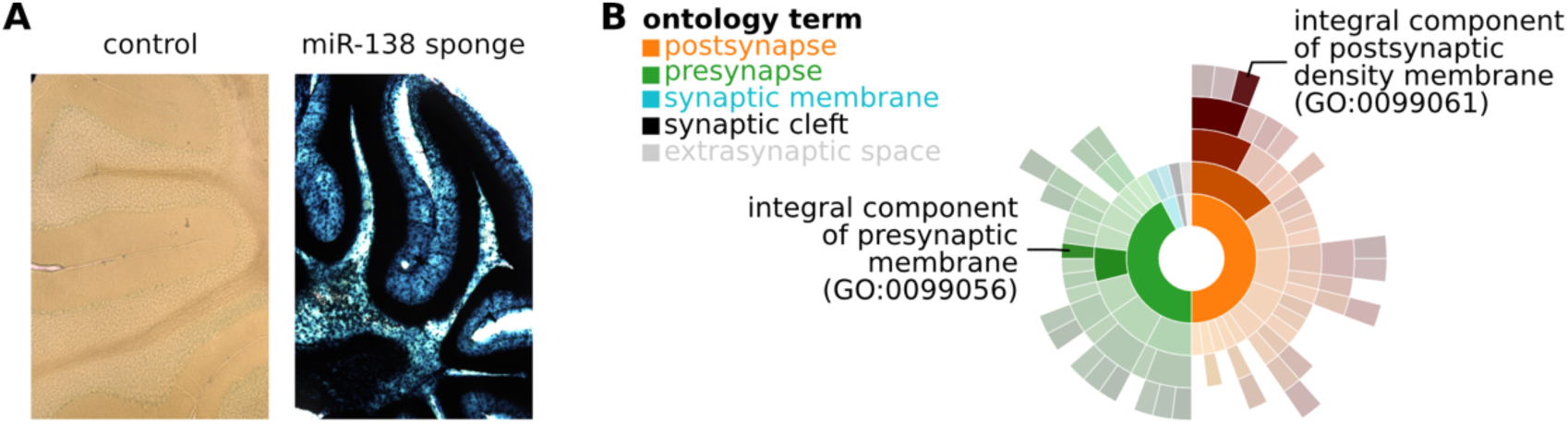
miR-138-5p sponge expression in the cerebellum and SynGO annotation of Erbb4. **(A)** lacZ staining shows strong expression of the 6×-miR-138 sponge in the cerebellar cortex of miR-138 sponge mice. Left: lacZ staining of a control mouse without CMV transgene (‘miR-138 floxed’, *Methods*). Right: lacZ staining of a miR-138 sponge mouse. **(B)** SynGO analysis of Erbb4 indicates presynaptic and postsynaptic localization.

**Figure S2.**
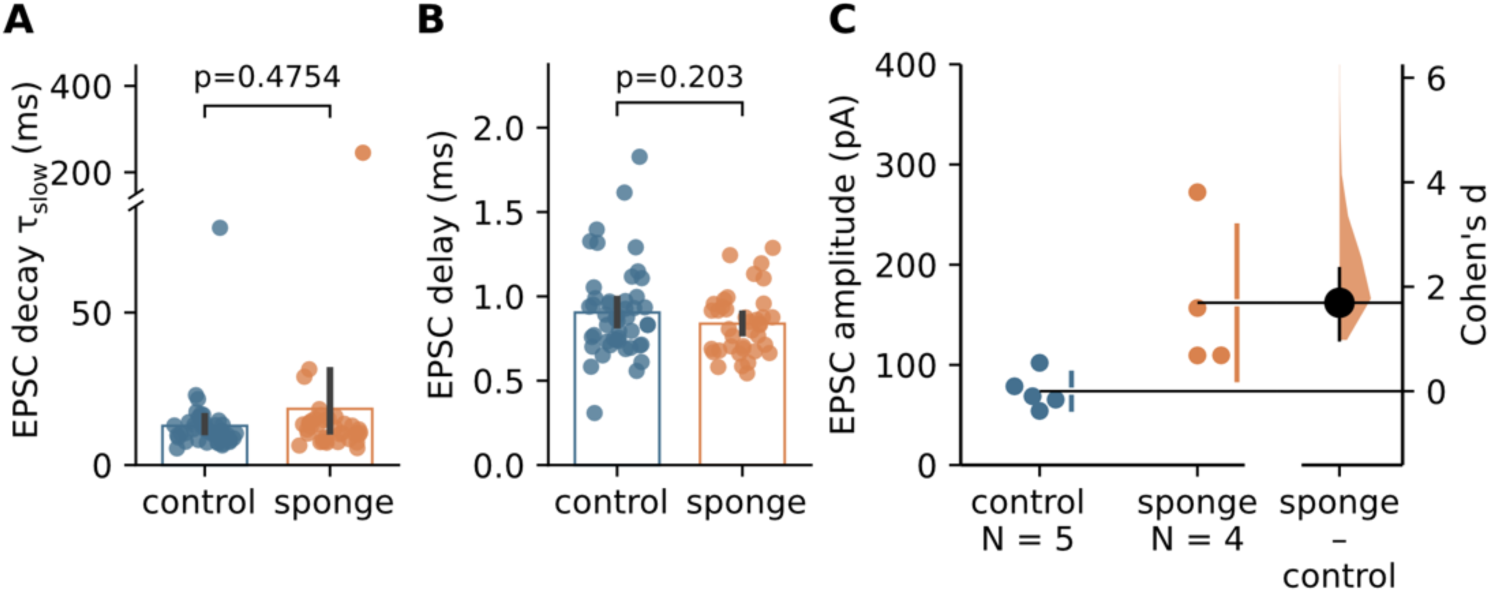
Additional EPSC quantification of MF–GC synapses. **(A)** Quantification of the slow EPSC decay time constant for control and sponge GCs. The slow EPSC decay component mainly reflects glutamate spillover from neighboring release sites. **(B)** EPSC delay for both genotypes, calculated as time between stimulation and onset of the EPSC. **(C)** EPSC amplitude quantification per animal for both groups, showing larger EPSCs in miR-138 sponge mice (Cohen’s d, 1.69 [95%CI 0.98, 2.35]).

**Figure S3.**
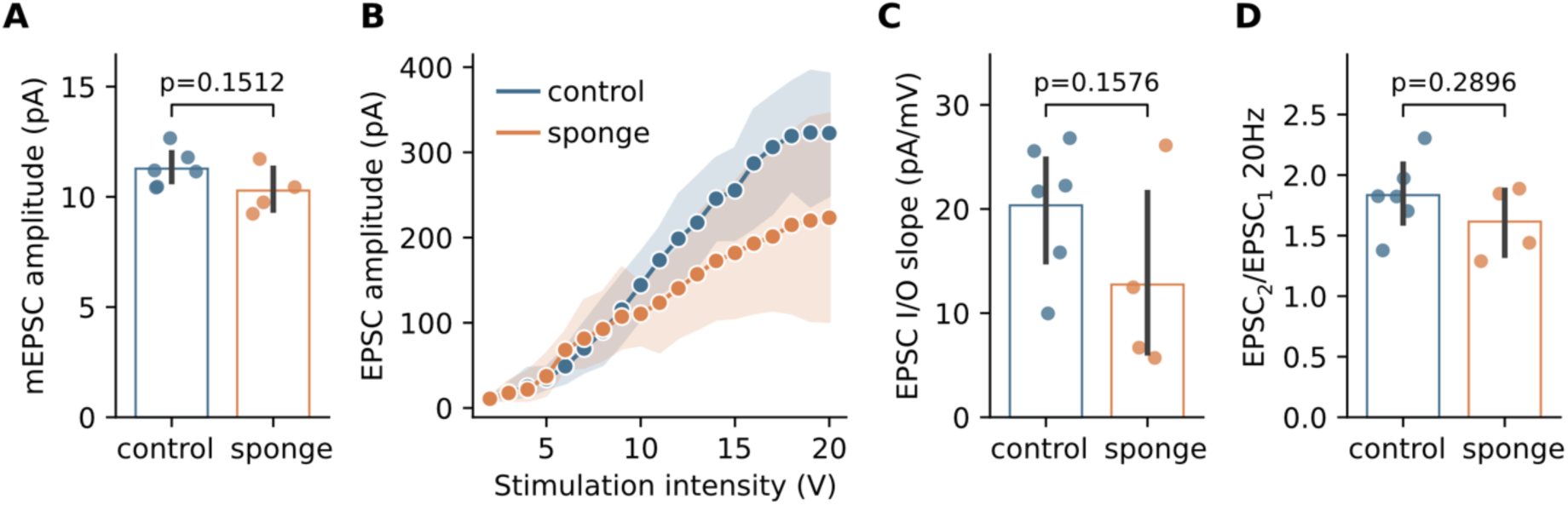
miR-138-5p sponge expression does not increase synaptic efficacy at PF–PC synapses. **(A)** Quantification of mEPSC amplitude for Purkinje cells of both genotypes, which was not increased in miR-138 sponge mice. **(B)** EPSC amplitude versus stimulation intensity for control and sponge parallel fiber to Purkinje cell (PF–PC) synapses. Dots are averages of ten EPSCs, shaded areas represent 95% CI. **(C)** Slopes of PF–PC EPSC input/output relationship obtained by linear regression. **(D)** Paired-pulse ratio (EPSC_2_/EPSC_1_) at 20 Hz stimulation frequency was similar in control and sponge PF–PC synapses.

**Figure S4.**
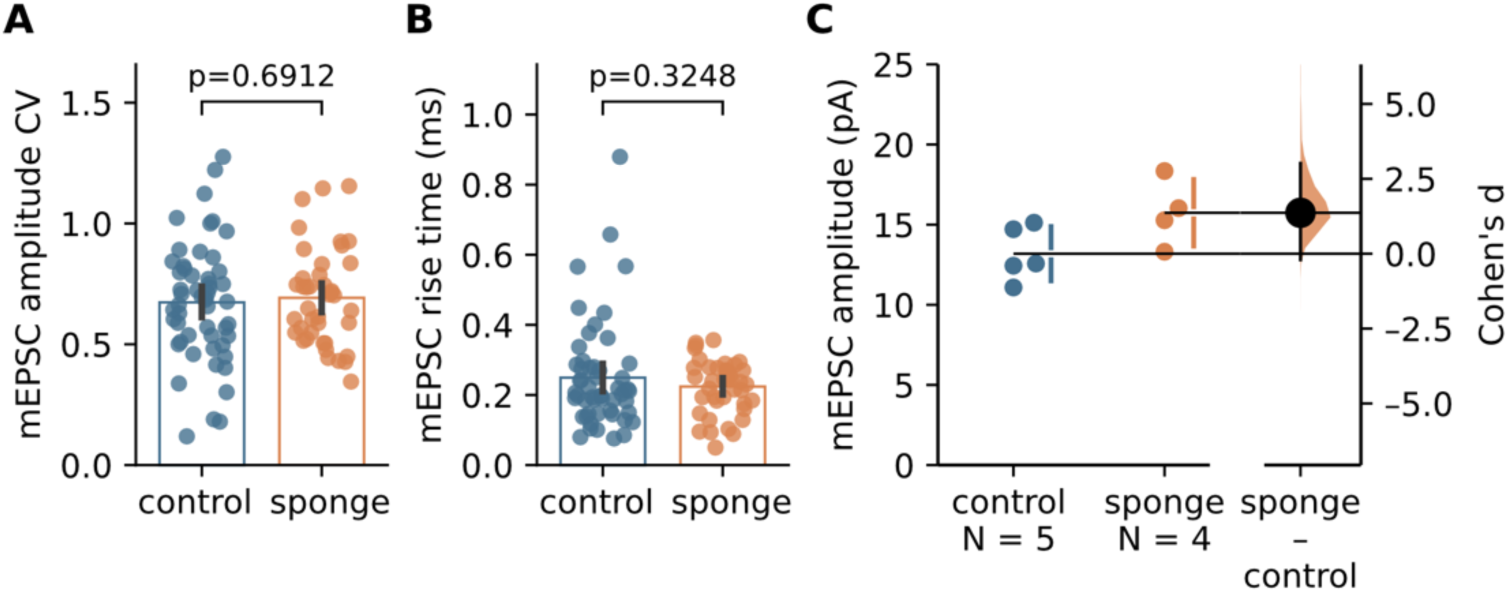
Additional mEPSC quantification of MF–GC synapses. **(A)** Coefficient of variation (CV) of mEPSC amplitudes for control and sponge GCs. **(B)** mEPSC 10–90% risetime for both genotypes. **(C)** mEPSC amplitude quantification per animal for both groups, showing larger mEPSCs in miR-138 sponge mice (Cohen’s d, 1.37 [95%CI −0.20, 3.02]).

**Figure S5.**
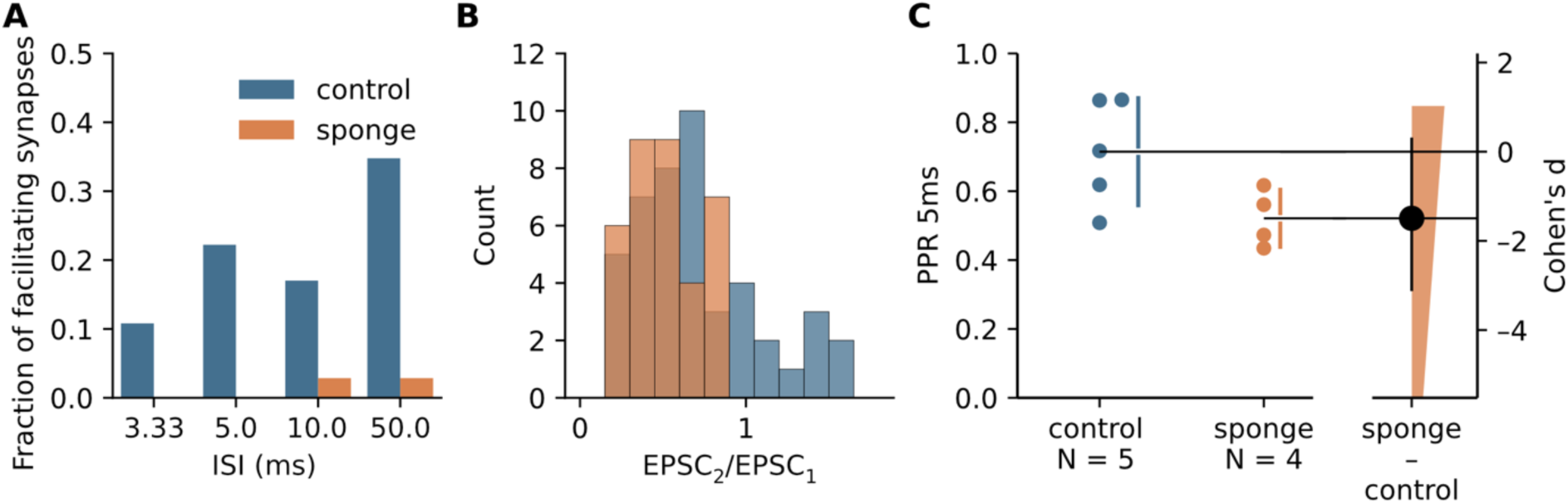
Absence of paired-pulse facilitation in miR-138 sponge GCs. **(A)** Fraction of MF–GC synapses showing facilitation (i.e., PPR >1) for different inter-stimulus intervals (ISIs). **(B)** Histogram of MF–GC PPRs across synapses at 5 ms ISI, showing a narrower distribution in miR-138 sponge mice. **(C)** PPR analysis on the level of individual animals. PPR (5 ms inter-stimulus interval) is reduced in miR-138 sponge GCs (Cohen’s d, −1.49 [95%CI −3.1, 0.29]), consistent with a change in release probability.

**Figure S6.**
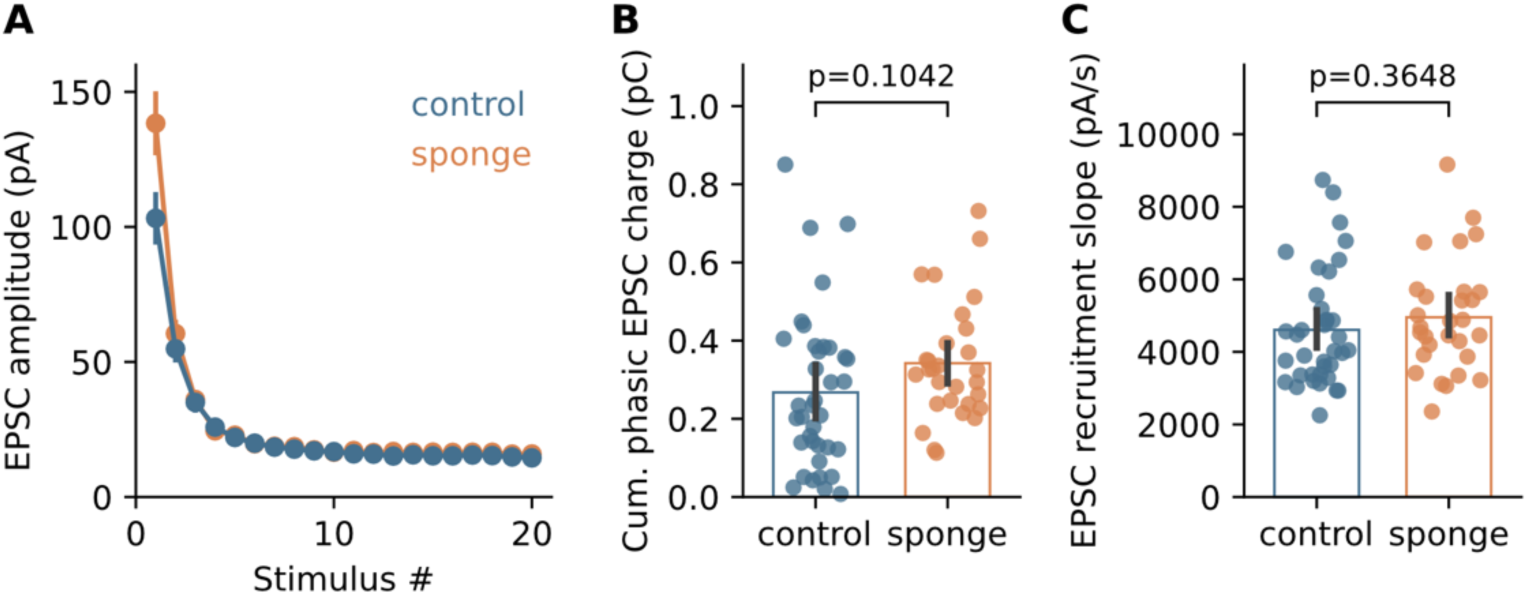
Additional analyses of 300-Hz train stimulation. **(A)** Absolute EPSC amplitude of 300-Hz train stimulation for control (blue) and miR-138 sponge (orange) GCs. **(B)** Cumulative phasic EPSC charge for both genotypes. **(C)** Recruitment slope for both genotypes, obtained by linear regression of the cumulative EPSC amplitude at steady state.

**Figure S7.**
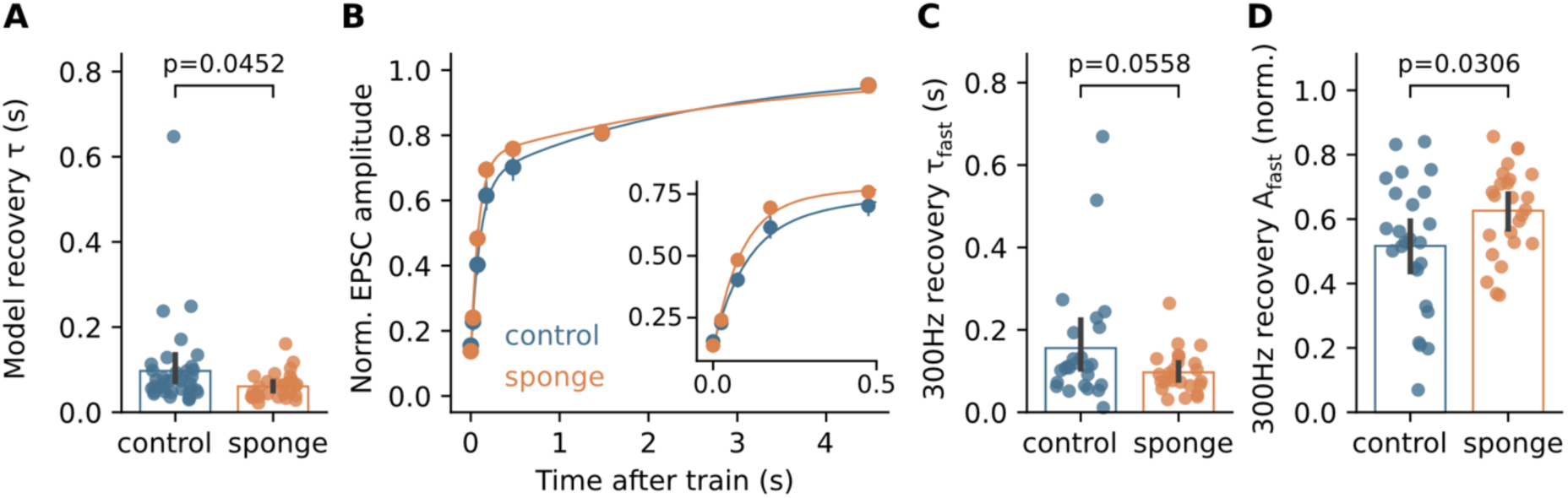
miR-138 sponge expression speeds recovery from synaptic depression. **(A)** The time constant of recovery from depression is faster in models fitted to sponge data from high-frequency train stimulation. **(B)** Normalized EPSC amplitude versus time after 300-Hz trains for both conditions. Lines are bi-exponential fits to the recovery time course. Inset shows initial phase of recovery on a shorter time scale. **(C)** The fast time constant of recovery from depression is shorter at miR-138 sponge synapses. **(D)** Quantification of the relative amplitude of the fast recovery component for both genotypes.

## Notes

### Competing Interest Statement

The authors have declared no competing interest.

